# The pyrethroid insecticide deltamethrin disrupts neuropeptide and monoamine signaling pathways in the gastrointestinal tract

**DOI:** 10.1101/2024.12.14.628386

**Authors:** Alexandria C. White, Ian N. Krout, Sabra Mouhi, Jianjun Chang, Sean D. Kelly, W. Michael Caudle, Timothy R. Sampson

## Abstract

Enteroendocrine cells (EECs) are a rare cell type of the intestinal epithelium. Various subtypes of EECs produce distinct repertoires of monoamines and neuropeptides which modulate intestinal motility and other physiologies. EECs also possess neuron-like properties, suggesting a potential vulnerability to ingested environmental neurotoxicants. One such group of toxicants are pyrethroids, a class of prevalent insecticides used residentially and agriculturally. Pyrethroids agonize voltage-gated sodium channels (VGSCs), inducing neuronal excitotoxicity, and affect the function of monoamine-producing neurons. Given their anatomical location at the interface with the environment and their expression of VGSCs, EECs likely represent a vulnerable cell-type to oral pyrethroid exposure. In this study, we used the EEC cell line, STC-1 cells, to evaluate the effects of the common pyrethroid deltamethrin on the functional status of EECs. We find that deltamethrin impacts both expression of serotonergic pathways and inhibits the adrenergic-evoked release of an EEC hormone, GLP-1, *in vitro*. In a mouse model of oral exposure, we found that deltamethrin induced an acute, yet transient, loss of intestinal motility, in both fed and fasted conditions. This constipation phenotype was accompanied by a significant decrease in peripheral serotonin production and an inhibition of nutrient-evoked intestinal hormone release. Together, these data demonstrate that deltamethrin alters monoaminergic signaling pathways in EECs and regulates intestinal motility. This work demonstrates a mechanistic link between pyrethroid exposure and intestinal impacts relevant to pyrethroid-associated diseases, including inflammatory bowel disease, neurodegenerative disease, and metabolic disorders.

---

Enteroendocrine cells (EECs) are sensory transducing cells of the gastrointestinal (GI) epithelium. Physically located at the interface of the intestinal-luminal environment, EECs are readily exposed to numerous factors, including intrinsic microbiome components and exogenous environmental and dietary molecules. In response to such stimuli, subsets of EECs produce various monoamines and neuropeptides including serotonin (5-hydroxytryptamine; 5-HT), CCK (cholecystokinin), PYY (peptide YY), and GLP-1 (glucagon-like peptide 1). These signaling molecules act to modulate intestinal motility, regulate blood glucose, and transmit satiety signals to the brain (1, 2). Although the role of EECs in microbial and nutrient sensing is well-described (3-7), whether and how EECs respond to common environmental toxicants is largely unknown.

Pyrethroids are a common insecticide used residentially (e.g., home pest control, gardening, and lice treatment), in veterinary medicine (e.g., flea and tick treatment), and in agricultural industries. Their prevalent and large-scale use in food production and agriculture results in widespread pyrethroid detection within foodstuffs and water, ultimately leading to chronic, low-dose oral exposures to the human population (8). In rare cases of acute toxicity, high-level exposure leads to neurological defects including seizures, discoordination and imbalance, and cognitive impairment (9). These central nervous system (CNS) impacts are likely due to pyrethroid actions on their main molecular targets, voltage-gated sodium channels (VGSCs). Pyrethroids are VGSC agonists, leading to their prolonged hyperactivation (10). While Type I pyrethroids act primarily on VGSCs, Type II pyrethroids simultaneously antagonize GABA_A_ receptors (GABARs), furthering neuronal excitation by preventing chloride-mediated inhibition. Through these mechanisms, experimental models have demonstrated that pyrethroid exposure leads to significant disruption of monoamine signaling in neurons, including serotonin and dopamine (11-15). However, the actions of pyrethroids on non-neuronal cells of the periphery are less described.

Given their location within the intestinal epithelium, EECs represent a likely first cellular target of pyrethroids and other ingested toxicants. Their potential vulnerability to pyrethroid toxicity is highlighted by their expression of pyrethroid-sensitive VGSCs. Using deltamethrin, a prevalent Type II pyrethroid insecticide, we assessed impacts to monoamine pathways and hormonal release in both an EEC culture model and in a murine oral exposure model. We find that deltamethrin dose-dependently interferes with expression of monoamine pathways in EECs *in vitro* and induces acute, but transient, intestinal dysmotility *in vivo*. Correspondingly, we observe that intestinal serotonin concentrations are significantly and transiently reduced after acute deltamethrin exposure. Lastly, we demonstrate that stimulated release of EEC neuropeptides is suppressed by acute deltamethrin exposure both *in vitro* and *in vivo* and is chronically associated with increased food intake. Overall, these data demonstrate a significant impact of a prevalent pyrethroid on EEC status and GI function, suggesting a mechanistic link for pyrethroid-associated diseases that impact the GI tract, including inflammatory bowel disease (16), metabolic syndrome (17-20), and Parkinson’s disease (21-23).

## Materials and methods

### Chemicals and treatments

For cell culture experiments, deltamethrin (Cat#: N-11579-250MG, purity 99%, Chem Service, Inc, Westchester, Pennsylvania) was dissolved in sterile dimethyl sulfoxide (DMSO, cell culture grade) to form a stock concentration of 100 mM that was aliquoted and stored at -20C until use. For mouse experiments, deltamethrin was dissolved in filter-sterilized corn oil (Mazola) and prepared fresh one day prior to each use for acute experiments or every 3 uses for chronic experiments, wherein it was stored in the dark at RT. Laboratory-grade acetone was used to facilitate deltamethrin’s dissolution and was allowed to evaporate overnight. For cell culture, glucose and epinephrine were both purchased from Sigma (Glucose Cat#: G8270-1KG, purity 99.5%; Epinephrine Cat#: E4250-1G, purity 99%), dissolved in PBS, and aliquoted for storage in -20C. Calcium-free media (Gibco Cat#: 21068028) was supplemented with 0.1mM EDTA, filter-sterilized, and stored at 4C. Ensure® (Abbott Pharmaceuticals) mixed-meal nutrient mixtures were prepared fresh on the day of use and mixed with 6% carmine red dye with 0.5% methylcellulose (Sigma Cat#: C1022 and M7027, respectively) prior to gavage. Unless stated otherwise, all stock solutions were filter-sterilized prior to use and were of pharmaceutical or food grade.

### Animal husbandry

C57BL/6J mice (The Jackson Laboratory, RRID: IMSR_JAX:000664) were co-housed according to treatment group in Emory University’s Whitehead Biomedical Research Building rodent facility. Mice were provided standard food and water *ad lib*, except when noted for mixed-meal nutrient stimulation experiments, under a 12h light/dark cycle with lighting at 7AM. At the experimental endpoints, 4-24h or 12 weeks following final treatments or exposures, mice were humanely euthanized via cardiac perfusion with sterile PBS after deep isoflurane anesthesia. All animal husbandry and experimental procedures were performed in accordance with Emory’s Institutional Animal Care and Use Committee (IACUC) protocol #201900030.

### Oral exposures

At 10-12 weeks of age, mice were orally gavaged once with 3 mg/kg deltamethrin (Chem Service, Cat#: N-11579-250MG) dissolved in filter-sterilized corn oil or filter-sterilized corn oil alone as vehicle control using a sterilized metal feeding needle (22G). This dose aligns with prior work (11, 24-26) and falls within previously reported no observable effect and no observable adverse effect limits defined by the U.S. EPA (27). For chronic exposure experiments, mice were gavaged once weekly for 12 weeks. Mice were monitored closely for overt signs of acute toxicity, none of which were observed throughout our studies. Subsequent intestinal and behavioral assays were conducted within consistent timeframes to account for circadian rhythm. For acute experiments, gavages occurred every 7 mins beginning at 10AM for 4h exposures or 3PM for 24h exposures, while carmine/mixed-meal nutrient gavages occurred every 7 mins starting at 12:30 PM, and fecal output and any motor behaviors occurred in the afternoons at 2PM. For chronic experiments, gavages occurred at 1PM once per week, while fecal output occurred the following morning at 10AM, and tests of total GI transit occurred at 0, 3, 6, and 12 weeks post-exposure and lasted from 10AM until 6PM. Food intake was measured only under chronic treatment conditions each week on the day prior to deltamethrin or vehicle gavage by weighing the amount of food consumed each week per cage, where food intake = amount of food added the prior week – amount of food remaining at the end of the week.

### Intestinal behaviors

#### Fecal output

Performed as previously described (28). At the indicated timepoints (Acute: 4h or 24h after single oral gavage; Chronic: 24h after each oral gavage), mice were placed in a sterile 1L plastic container to count the number of fecal pellets produced every 5 mins for 30 mins.

#### Intestinal motility assay

Performed as described previously (29). At the indicated timepoints, mice were orally gavaged with 100 μL 6% carmine red dye 1.5h prior to sacrifice to visually track motility. During tissue collection, the distance traveled (cm) by the carmine dye was recorded as a percentage of the total length of the gastrointestinal tract (small intestine length + large intestine length, excluding cecum). If the dye was in the cecum, it was recorded as traveling 100% of the small intestine length. In a second set of experiments, Ensure (Abbott Pharmaceuticals) was pre-mixed with 6% carmine red dye prior to each mouse receiving 100 μL of the mixture 1.5h prior to sacrifice.

#### Total intestinal transit assay

Performed as previously described in detail (30). After one hour of habituation in an isolated behavioral testing room, mice were orally gavaged with 100 μL carmine red dye solution [6% carmine red dye mixed with 0.5% methylcellulose]. One hour later, mice were split into separate cages where they were housed individually with access to food and water but no bedding. Mice were observed every 15 mins for up to 8 hours or until a red fecal pellet was produced, at which point mice were returned to their home cage. The difference in time at which the mouse is gavaged and a red pellet is produced equals total intestinal transit time and was recorded in hours.

### Cell culture

The mouse (*Mus musculus*) secretin tumor cell line (STC-1) at passage 29 was purchased from the American Type Culture Collection (ATCC, Cat#: CRL-3254) and cultured as described in detail (31). Cells were seeded in 24 or 96-well plates at a density of 80,000 or 20,000 cells/well, respectively. Cells were cultured in DMEM/F12 media with GlutaMAX (GIBCO, Cat#: 10565018) supplemented with 10% heat-inactivated fetal bovine serum (FBS; Gibco, Cat#: 16140071) and 2% penicillin/streptomycin (5,000 U/mL, Gibco, Cat#: 15070063), modified from work described previously (32). Cells were incubated in a 5% CO_2_ humidified chamber held at 37C, and passage numbers were maintained between 30 and 35. Cells were passaged via 5-min incubation in 5mL 0.25% Trypsin-EDTA (Gibco, Cat#: 25200072). Media was changed every other day until treatments, which occurred 3-4 days after seeding once cells reached ∼80% confluency.

### Cytotoxicity Assays

Chemical-mediated cell death was assessed using the colorimetric CytoTox 96® Non-Radioactive Cytotoxicity Assay (Promega, Cat#: G1780), which measures lactate dehydrogenase (LDH) released into the supernatant, according to manufacturer’s instructions and described in detail previously (33). LDH was quantified either 1h (for GLP-1 release experiments) or 24h (for gene expression) after treatments. Cells were grown to confluency (approximately 48-72h after seeding at 10k cells/well in a 96 well plate) and maintained at 37C until treatments. Maximum LDH release was measured by adding 10 μL lysis buffer per 100 μL media to a subset of untreated control wells and allowing them to incubate for 45 mins. Afterward, 50 μL of supernatant from each well was collected and placed in a new 96 well plate for LDH quantification along with 50 μL of CytoTox 96 Reagent and incubated for 30 mins at room temperature (RT) in the dark. Stop solution was added immediately prior to absorbance reading, which was measured at 490nm using a spectrophotometer.

### Gene expression analysis

#### RNA extraction

At confluency, STC-1 cells were treated with 0.01% dimethyl sulfoxide (DMSO) vehicle control or 12.5, 25, 50, or 100 μM deltamethrin. Approximately 24h later, cells were collected for RNA extraction as described in detail previously (34). Briefly, cells were washed gently once with PBS and 250 μL Trizol (Zymo Research, Cat#: R2050-1-200) was added to each well for RNA extraction using the Qiagen RNeasy Mini Kit (Cat#: 74106). In brief, cell lysates were homogenized by sonication (QSonica, ∼35-45 setting) for 3-5s before 1/5 volume of chloroform was added to each sample and shaken for 15s. After sitting for 2 mins at RT, samples were centrifuged at 12,000 rpm for 15 mins at 4C. The upper aqueous clear layer was transferred to a QIAshredder (Qiagen, Cat#: 79656) and centrifuged for 2 mins at full speed at RT. The flow-through was combined with a half volume of 70% ethanol and transferred to an RNeasy spin column. Samples were centrifuged briefly for 30s at 10,000 rpm and flow-through was discarded. 700 μL Buffer RW1 was added to the spin column prior to centrifuging again for 30s at 10,000 rpm. Flow-through was discarded and 500 μL Buffer RPE was added to the spin column followed by centrifuge for 30s at 10,000 rpm. This step was repeated but instead centrifuged for 2 mins before discarding flow-through. The RNeasy spin column was then placed in a new collection tube and centrifuged for 1 min at full speed to collect any excess liquid. Finally, the spin column was placed in a new 1.5 mL Eppendorf tube, 30 μL RNase-free water was added, and samples were centrifuged at 10,000 rpm for 1 min to elute RNA. RNA concentrations for each sample were measured using a NanoDrop.

#### cDNA conversion

RNA was converted to cDNA using the iScript cDNA Synthesis Kit (Bio-Rad, Cat#: 1708891) following the manufacturer’s instructions and described in detail previously (35). Briefly, samples were diluted to 50 ng/μL up to a total of 1 μg RNA. To achieve a 20 μL reaction volume, 4 μL of 5x iScript Reaction Mix and 1 μL iScript Reverse Transcriptase was added for every 15 μL of sample. The complete reaction mix was then incubated in a thermal cycler as follows: priming for 5 mins at 25C, reverse transcription for 20 mins at 46C, inactivation for 1 min at 95C, and an optional hold step at 4C.

#### Real-time quantitative PCR (RT-qPCR)

was performed for gene expression analysis according to a previously described protocol (36). cDNA was diluted 1:10 to reach a final amount of 10 ng cDNA per reaction. Primer pairs were diluted from 100 μM stocks to 10 μM combined working stocks. Per sample and gene of interest, a 12 μL reaction mix was created: 6 μL SYBR Green PCR Master Mix (Thermo Fisher Scientific, Cat#: 4309155), 2 μL diluted primer pairs, 2 μL DI H_2_O, and 2 μL diluted cDNA. Each sample was tested in technical duplicate and appropriate controls and blanks were used (no-template control, cDNA only, DI H_2_O only). All primer pair sequences are reported in Supplementary Table 1.

### Protein quantifications

#### ELISAs

Protein preparation and western blotting was performed as described in detail previously (37). Indicated cell suspensions were homogenized via sonication (QSonica, ∼35-45 setting, 3-5s per sample) in ice-cold Meso-Scale Discovery (MSD) homogenization buffer (1.0 M Tris, 0.5 M MgCl2, 0.1 M EDTA, deionized water, and 1% Triton-X 100, plus added protease and phosphatase inhibitors [Thermo Scientific, Cat#: A32961]). Lysed samples were then centrifuged at 4C for 10 mins at 20,000x*g* and the protein-rich supernatant was collected and stored at -80C until analysis. Protein levels were quantified using a standard Pierce BCA protein assay kit (Thermo Scientific, Cat#: A55864). ELISAs were performed according to manufacturer’s instructions and described in detail previously (38). Active GLP-1 release in STC-1 cell supernatant was detected with a GLP-1 (Active) ELISA kit (Millipore, Cat#: EGLP-35K). Samples were diluted 10-fold into kit-provided Assay Buffer prior to adding 200 μl sample/well. The plate was incubated overnight at 4C followed by five washes prior to the addition of 200 μl Detection Conjugate which incubated on a slow shaker for 2 hours at RT. The plate was washed 3 times and Substrate was added to each well for a final 20-min incubation in the dark at RT. Once sufficient fluorochrome was generated, 50 μl Stop Solution was added before reading the plate at an excitation/emission wavelength of 355 nm/460 nm.

#### Multiplexed ELISAs

Used to detect gut hormones in serum, blood derived from cardiac puncture was phase separated into serum via Vacuette tubes (Greiner-Bio One, Cat#: 454243P) after centrifugation at RT for 10 mins at 1800 rcf and stored at −80C until sample analysis. Multiplexed ELISAs were performed on indicated serum samples through the Emory University Multiplexed Immunoassay Core. Experiments were performed in duplicate using the U-PLEX Metabolic Hormones Combo 1 for mouse (Meso Scale Discovery, Rockville, MD, USA, Cat#: K15306K-2) with ‘blank’ replicates that served as negative controls, according to the manufacturer’s instructions. All samples were diluted two-fold for analyses.

### High Performance Liquid Chromatography (HPLC)

HPLC was performed on indicated samples (ileum, colon, serum) through the Emory University HPLC Bioanalytical Core. Samples were resuspended in 300 µL ice cold 0.1M PCA, 0.1 mM EDTA and sonicated on dry ice using probe sonication. Pulses were 1s on, 10s off, and used an amplitude of 25% for 20-30s (QSONIC Q500A, Newtown CT). The homogenated samples were then centrifuged at 13,000 × g for 15 mins at 4C. Sample supernatants were transferred into new 0.22 µM PVDF microcentrifuge filter tubes and filtered through a spin filter at 5000 × rpm for 5 mins at 4C. Reverse-phase HPLC with electrochemical detection was used to measure monoamine concentrations. Protein pellets were dissolved in 500 µL 2% Sodium dodecyl sulfate (SDS). Protein quantification was performed in triplicate in 96-well microplates with SpectraMax M5e spectrophotometer (Molecular Devices, Sunnyvale, CA) using the Pierce BCA Protein Assay Kit (Thermo Scientific, Cat #A55864). Monoamines were quantified using the ACQUITY ARC system equipped with a 3465 electrochemical detector (Waters). Separations were performed using an Xbridge BEH C18, 2.5 µm, 3 x 150 mm column (Waters) at 37C. The mobile phase contained 100 mM citric acid, 100 mM phosphoric acid, pH 3.3, 0.1 mM EDTA, 525 mg/L 1-octanesulfonic acid, and 7% acetonitrile. The detection flow cell was SenCell with 2 mm GC WE and the cell potential was set at 800 mV with a salt bridge reference electrode. The AST position was set at 1 while the ADF was 0.5 Hz. The needle was washed with water and the pump piston was washed with 15% isopropanol. 20 µL of each sample was injected before being eluted isocratically at 0.6 mL/min. The analytes were identified by matching criteria from retention time measures to known standards (Sigma Chemical Co., St. Louis MO). Compounds were quantified by comparing peak areas to those of standards.

### Statistical Analyses

All datasets were analyzed using GraphPad Prism 9 statistical software. Comparisons between two groups were generated using two-tailed t-tests for normally distributed data or by Mann Whitney nonparametric tests for non-normally distributed data. Comparisons between more than two groups were analyzed with one-way (one independent variable) or two-way (two independent variables) ANOVAs and follow-up post-hoc tests. Dunnett’s multiple comparisons test was used to compare means to a control mean, Šídák’s multiple comparisons test was used to compare pre-selected means to one another, and Tukey’s multiple comparisons test was used to compare all means to one another. Where indicated, multiple t-test comparisons were performed. Outliers were removed according to the ROUT outlier identification test. All numerical data and statistical outputs are included as a supplemental data file.

## Results

### Deltamethrin dysregulates monoamine pathways in EECs

To address the effects of deltamethrin on EECs, we first used the STC-1 cell line, a murine transformed cell line that is broadly representative of EEC subtypes (32). We observe that this line expresses a set of VGSCs, namely *Scn2a-Scn11a*, excluding *Scn5a* (Supplementary Fig. S1A). Other than *Scn2a* and *Scn9a*, these VGSCs are sensitive to deltamethrin and other pyrethroids (39). Over a range of relevant doses (0-100µM), we observed little cytotoxicity compared to vehicle controls following a 24hr deltamethrin exposure (Supplementary Fig. S1B). VGSC expression, specifically *Scn2a, Scn4a,* and *Scn9a* (but not *Scn3a, Scn5a, Scn8a, Scn10a,* or *Scn11a*), is greatly increased at higher doses at this timepoint (Supplementary Fig. S1C), suggesting that these channels are indeed affected following deltamethrin exposure.

Since deltamethrin disrupts monoaminergic signaling in CNS neurons (11, 13, 14, 40), we next sought to determine whether these pathways were similarly affected within these specific intestinal endocrine cells. Targeted gene expression analysis following 24hrs of deltamethrin exposure revealed that deltamethrin significantly dysregulates components of monoamine function, in particular serotonin synthesis and release pathways that are central to EEC functions. This includes significant increases in *Tph2*, *Vmat2*, *Slc6a4*, *MaoA*, and *Comt* expression and decreases in *Tph1* but no effects on *Vmat1* or *Ddc* (Fig. 1A). Thus, deltamethrin dysregulates monoamine pathways in EECs.

**Figure 1.**
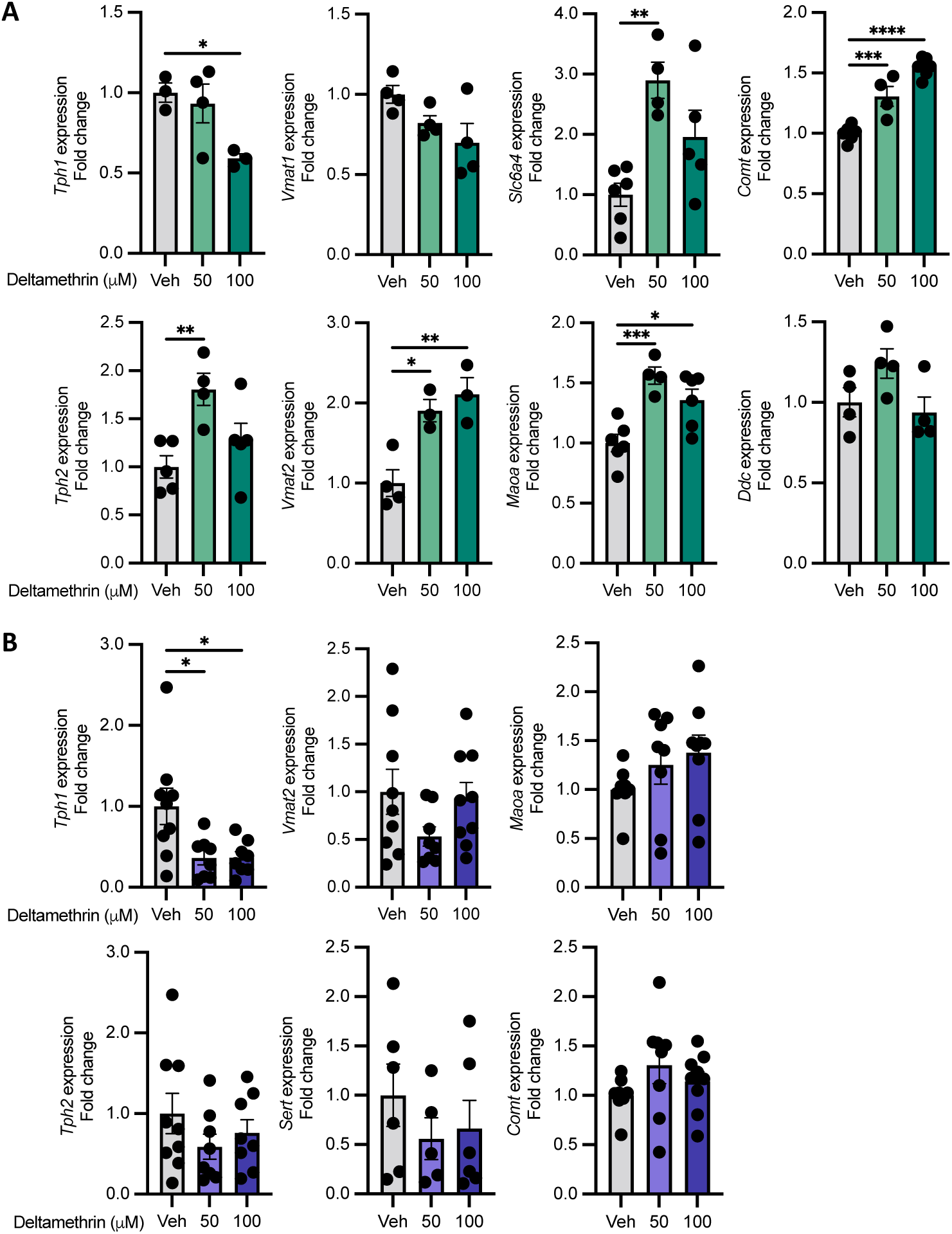
Deltamethrin dysregulates monoaminergic gene expression pathways in STC-1 cells in a calcium-dependent manner. **A** Relative expression values expressed as fold change from STC-1 cells treated with DMSO vehicle alone (Veh), 50, or 100 µM deltamethrin for 24h**. B** Relative expression values expressed as fold change from STC-1 cells treated with DMSO vehicle alone (Veh), 50, or 100 µM deltamethrin for 24h in calcium-free media. N = 3-6 (**A**) or n = 6-9 (**B**) independent culture wells per condition. Points represent independent wells (means of technical duplicates) and bars the mean ± SEM. Data compared by ordinary one-way ANOVA with Dunnett’s multiple comparisons test comparing each group to the control mean. \**p*≤0.05, \*\**p*≤0.005, \*\*\**p*≤0.001, \*\*\*\**p*<0.0001.

VGSC activation, by pyrethroids and other agonists, leads to an influx of calcium to promote downstream signaling cascades (10). In order to determine whether the impacts we observed on gene expression in the monoamine pathways were dependent on calcium signaling, we performed identical deltamethrin exposures in the absence of extracellular calcium. Unlike deltamethrin exposures in calcium-replete media, in the absence of calcium, deltamethrin exposures did not result in the same widespread dysregulation of monoamine pathway genes (Fig. 1B). Excluding *Tph1*, expression of other genes, such as *Tph2, Vmat2*, *Slc6a4*, *Maoa,* and *Comt* was unaffected. Therefore, the ability of deltamethrin to dysregulate these serotonergic and monoaminergic pathways in EECs is dependent on calcium signaling, and likely their canonical activation of VGSCs.

### Deltamethrin inhibits epinephrine-evoked GLP-1 release in STC-1 cells

Other than monoamines, EECs produce a variety of neuropeptide hormones, including GLP-1, among others (32). Given that STC-1 cells robustly produce GLP-1 in response to various stimuli (41), we first tested whether deltamethrin impacted GLP-1 gene expression pathways. We found that GLP-1 receptor (*Glp1r*) expression is increased by deltamethrin in a calcium-dependent manner (Fig. 2B, C). We then assessed how deltamethrin functionally impacts EEC neuropeptide signaling by measuring GLP-1 release. STC-1 cells were treated with epinephrine or glucose, two compounds known to stimulate GLP-1 release in these cells (32, 42), resulting in a dose-dependent increase in GLP-1 release, as anticipated (Fig. S2A, B). To determine how deltamethrin may affect the ability of EECs to respond to these stimuli, cells were pre-treated with deltamethrin prior to an epinephrine stimulus. While deltamethrin alone did not affect GLP-1 release (Fig. 2D), pre-treatment with deltamethrin significantly inhibited epinephrine-mediated GLP-1 release (Fig. 2D). These data indicate that deltamethrin interferes with the ability of EECs to respond to key modulatory signals, resulting in significant functional dysregulation across monoamine and GLP-1 signaling pathways.

**Fig. 2.**
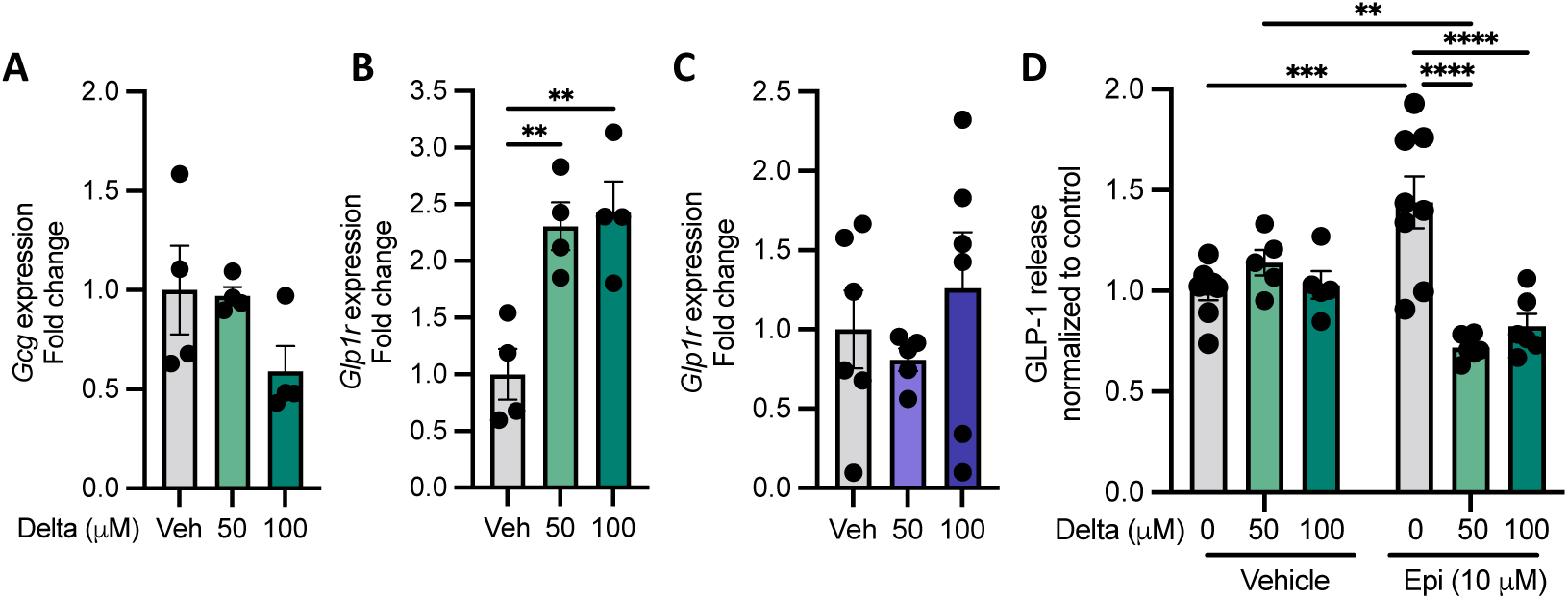
Deltamethrin inhibits epinephrine-induced GLP-1 release in STC-1 cells. **A-B** Gene expression of *Gcg* and *Glp1r,* respectively, normalized to control values after treatment with 0, 50, or 100 µM deltamethrin. **C** *Glp1r* expression after treatment with 0, 50, or 100 µM deltamethrin in calcium-free media. **D** GLP-1 release quantified by ELISA and normalized to control after 0, 50 or 100 µM deltamethrin treatment in the presence and absence of 10 µM epinephrine. **A-D** All data points represent averages of technical duplicates from individual samples where n = 4 (**A, B**), n = 5-6 (**C**), or n = 5-8 (**D**) per group. Data are depicted as mean ± SEM and compared by ordinary one-way ANOVA with Dunnett’s multiple comparisons test comparing each group to the control mean (**A-C**) or ordinary two-way ANOVA with Tukey’s multiple comparisons test comparing means all means to one another (**D**). \*\**p*<0.005, \*\*\**p*<0.001, \*\*\*\**p*<0.0001.

### Oral deltamethrin exposure induces acute constipation in mice

Given the significant impact on motility-regulating monoamine and GLP-1 pathways we observed following deltamethrin treatment of EEC-like cells in culture, we sought to determine potential intestinal functional impacts *in vivo*. We performed a battery of GI assessments in mice following an oral exposure to low-dose deltamethrin (3 mg/kg body weight). After an acute exposure, 4hrs post oral gavage (serum C_max_ time), we observed that deltamethrin-treated mice displayed significantly reduced fecal pellet output, compared to animals treated with corn oil vehicle (Fig. 3A). Moreover, a terminal dye transit assay demonstrated that intestinal motility was significantly delayed after deltamethrin exposure (Fig. 3B). By 24hrs post-exposure, fecal output in treated animals was restored and intestinal transit was improved albeit still impaired compared to vehicle controls, suggesting that deltamethrin’s effects, while drastic, are transient (Fig. 3C, D). These data clearly demonstrate that a single, low-dose, orally-administered deltamethrin exposure is sufficient to induce a constipation-like phenotype in mice.

**Figure 3.**
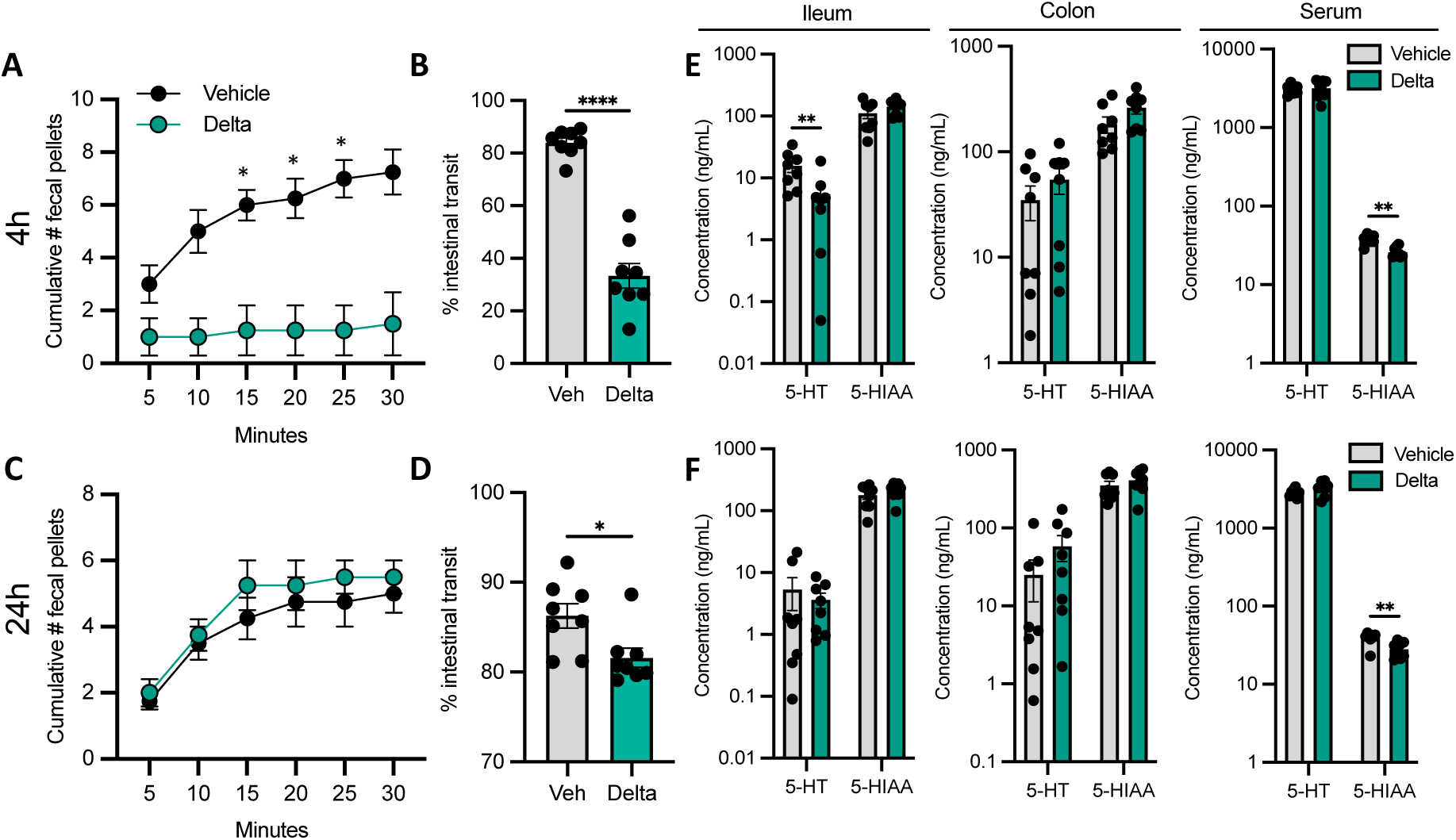
Oral deltamethrin exposure induces acute intestinal dysmotility and limits serotonin production. **A** Fecal output (# pellets produced in 30 mins) after 4h deltamethrin or vehicle exposure. **B** The % intestinal transit of red carmine dye in the GI tract of 4h vehicle or deltamethrin-treated mice immediately after sacrifice. **C** Fecal output after 24h deltamethrin or vehicle exposure. **D** The % intestinal transit of 24h vehicle or deltamethrin-treated mice. **E-F** Peripheral serotonin (5-HT) and 5-HIAA levels at 4h (**E**) or 24h (**F**) post vehicle or deltamethrin exposure in ileum, colon, and serum, as measured by HPLC. All data points represent averages of biological (**A, C**) or technical (**B, D-F**) replicates where n = 4 (**A, C**) or n = 8 (**B, D-F**) per group. Data are depicted as mean ± SEM and compared by ordinary Two-way ANOVA with Šídák’s multiple comparisons test between treatment groups (**A, C**), Mann Whitney test for non-normally distributed data (**D, E, F**), or two-tailed unpaired student’s t-tests (**B, E, F**). \**p*<0.05, \*\**p*<0.005, \*\*\*\**p*<0.0001.

Since we observed a striking *in vivo* intestinal motility phenotype that correlated with dysregulation of monoamine pathways in EECs in culture, we next measured circulating and tissue-resident serotonin levels. We detected significantly lower concentrations of serotonin (5-HT) in the ileum and lower 5-hydroxyindoleacetic acid (5-HIAA, the major serotonin metabolite) levels in serum, with no difference in colonic concentrations at acute (4hrs) timepoints post-exposure (Fig. 3E). At 24h post-exposure, we observed only a significant reduction in serum 5-HIAA, with no differences between treatment groups in the ileum or colon (Fig. 3F). These data together demonstrate that oral deltamethrin exposure significantly and acutely disrupts intestinal motility and accompanying peripheral monoamine signaling pathways.

### Deltamethrin interferes with nutrient-stimulated intestinal hormone signaling *in vivo*

GLP-1 is released *in vivo* in response to glucose and other nutritional stimuli (41). Given our observations that deltamethrin inhibits GLP-1 release in EECs *in vitro*, we assessed how deltamethrin modulates intestinal hormones, including GLP-1, in response to a high nutrient, mixed-meal stimulus *in vivo*. Mice treated with either deltamethrin or a corn oil vehicle were fasted for 4-6h and stimulated with a mixed-meal nutrient oral bolus (Fig. 4A). Similar to our prior experiment, oral deltamethrin exposure resulted in an acute, but transient intestinal dysmotility compared to vehicle controls, even in the presence of a mixed-meal (Fig. 4B, D). To determine the extent to which deltamethrin may interfere with gastrointestinal hormone signaling *in vivo,* we measured an array of circulating intestinal neuropeptide hormones. While GLP-1 concentrations themselves were not affected *in vivo*, we observed significantly less circulating insulin and leptin following acute deltamethrin exposure (Fig. 4C). This effect was transient, with concentrations similar to vehicle controls 24h post-exposure (Fig. 4E). Since both insulin and leptin contribute to satiety, and deltamethrin inhibited their release following a mixed meal, we tested whether deltamethrin modified food intake. Using a chronic treatment paradigm that allows the capture of weekly food intake, we observe an increase in food intake in deltamethrin-exposed mice (Fig. 4F, G), as well as qualitatively impaired GI transit (Fig. 4H). As a whole, these data emphasize a role for deltamethrin in perturbing broad gastrointestinal functions through neuropeptide modulation.

**Figure 4.**
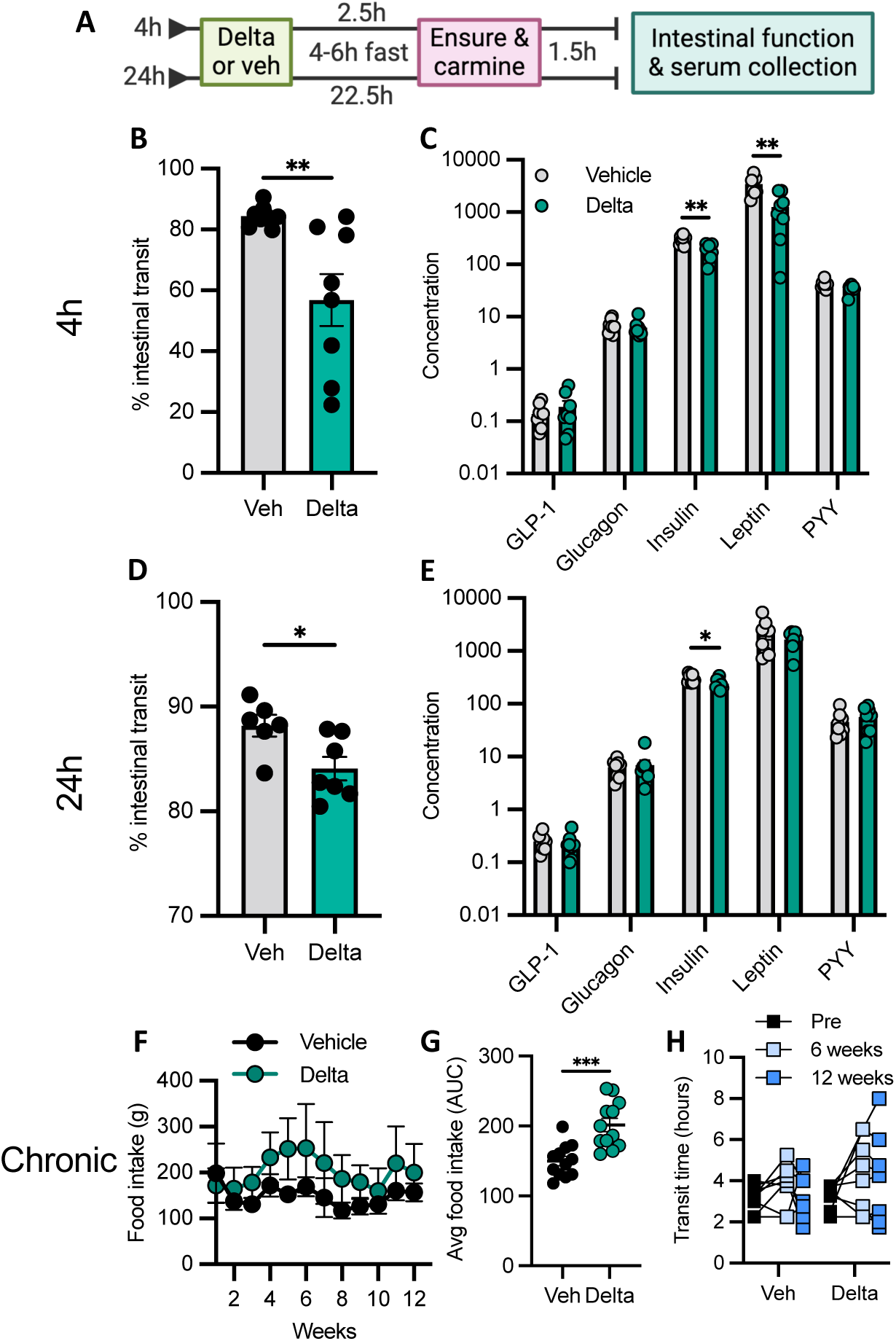
Deltamethrin inhibits release of intestinal hormones following a mixed-meal nutrient-stimulus. **A** Schematic representation of the experimental timeline. **B** % intestinal transit of carmine red dye 4h after exposure to either vehicle or deltamethrin and measured upon sacrifice. **C** Accompanying gut hormone panel in serum after 4h exposure to vehicle or deltamethrin, determined by multiplexed ELISA. **D** % intestinal transit of carmine dye after 24h exposure to vehicle or deltamethrin. **E** Corresponding quantification of gut hormone levels in serum after 24h exposure to vehicle or deltamethrin. **F** Average weekly food intake after chronic exposure to vehicle (black) or deltamethrin (green). **G** Total intestinal transit time in vehicle or deltamethrin-treated mice after 0 (black), 6 (light blue), or 12 weeks (blue) of exposure. **H** Average food intake expressed as area under the curve for vehicle- and deltamethrin-treated mice. **C, E** Concentration units are in pM for GLP-1 (active) and glucagon, uIU/mL for insulin, and pg/mL for leptin and PYY (total). **B-H** All data points represent individual mice (**B, D, H**), their averages after technical duplicates (**C, E**), or the average of two cages of 4 mice per cage (**F, G**), where n = 8 per group. Bars are depicted as mean ± SEM and compared by unpaired student’s t-test (**B, C, E, G**), Mann Whitney nonparametric tests for non-normally distributed data (**C, D**), or a two-way repeated measures ANOVA (**F, H**). \**p*<0.05, \*\**p*<0.005, \*\*\**p*<0.0005.

## Discussion

Enteroendocrine cells are among the first cell types to interact with ingested environmental toxicants. Their anatomical location at the interface between the intestinal luminal environment and the nervous system, in conjunction with their neuron-like properties, underscore their vulnerability to neurotoxic compounds. Because pyrethroids are highly prevalent and a common route of their exposure is ingestion (8), they are a relevant exposure to impact EEC function and intestinal physiology. Here, we show that the pyrethroid deltamethrin directly affects monoaminergic properties in EECs *in vitro* and induces *in vivo* perturbations to the gastrointestinal system that are associated with dysregulation of EEC functions, including serotoninergic signaling and neuropeptide hormone release. These findings not only establish EECs as vulnerable to pyrethroid toxicity, but also provide insight into the mechanisms by which deltamethrin alters intestinal physiology, which may underlie known gastrointestinal dysfunctions of exposure-associated diseases.

Several studies report various GI disturbances following high-dose or chronic pyrethroid exposures in both animals and humans (8, 9, 16, 43, 44). These include nausea, intestinal pain, dysmotility, and gastrointestinal inflammation—pathologies that overlap with clinical features of irritable bowel syndrome (IBS), which is associated with pyrethroid exposure (45), and inflammatory bowel disease (IBD) (16). As dysregulation of both serotonin and neuropeptide signaling contribute to these diseases and pathologies (46-50), our data supports a direct pathological contribution of deltamethrin exposure to the EEC dysfunctions that characterize these intestinal diseases.

We observe pyrethroid-induced suppression of physiologically stimulated GI hormones GLP-1, insulin, and leptin, which each have broad actions in GI physiology and metabolism (51-53). Pyrethroid exposure is linked to impairments in glucose metabolism, including diabetes (18, 19, 54, 55), with mixed findings to obesity (56, 57). Interestingly, serotonin has also been implicated as a contributor to obesity and diabetes (58). Given our observations that deltamethrin directly impacts both neuropeptide hormones and serotonin pathways in EECs, it suggests that EEC dysfunctions may be a link between pyrethroid toxicity and further metabolic disruption in the GI tract and systemically.

More broadly, pyrethroid exposure is linked with varying degrees of strength to neurodevelopmental disorders such as autism and attention-deficit hyperactivity disorder (13, 24, 59) and neurodegenerative outcomes relevant to both Parkinson’s disease (PD) (11, 14, 22, 60) and amyotrophic lateral sclerosis (ALS) (61), many of which involve established disruptions to monoaminergic pathways, including serotonin (62, 63). PD in particular presents with significant GI dysfunctions (64), though the specific intestinal pathologies which underlie these are not well described. Constipation is a prevalent prodrome of PD (65-68). It is tempting to speculate that environmental exposures, such as to pyrethroids, trigger early pathologies at the site of exposure within the GI tract. Given the evidence that pyrethroids impact monoamine signaling in the CNS, including serotonin signaling (12, 15, 69), it is plausible that these insecticides contribute to GI dysfunctions through similar mechanisms. Indeed, our data herein support the notion that pyrethroid exposures are sufficient to induce dysfunction within vulnerable intestinal cells that lead to relevant prodromal features of these neurological diseases.

This study directly tests pyrethroid-monoamine interactions within the GI tract, a critical step toward understanding the broad health impacts of pyrethroid exposure on intestinal physiology that is linked to many pyrethroid-associated diseases. We provide compelling evidence that the pyrethroid deltamethrin alters intestinal pathways important for serotonin trafficking in EECs, acutely impairs intestinal motility, and diminishes intestinal hormone production in response to physiological stimuli. Our data highlight the continued need to study toxicological impacts in the GI tract that may be associated with long-term exposure to these prevalent chemicals and underlie those diseases epidemiologically linked to exposure.

## Data availability

All numerical data and statistical outputs are available in the supplemental raw data file associated with this manuscript.

## Acknowledgments

We thank Isabel Fraccaroli, Tiffany Chen, and Shreya Bandlamudi for scientific support; and members of the Emory Division of Animal Resources for technical support. We acknowledge support from the Emory Multiplexed Immunoassay Core (EMIC) and the Emory HPLC Core which are subsidized by the Emory University School of Medicine as Integrated Core Facilities and are supported by the Georgia Clinical and Translational Science Alliance of the NIH (UL1TR002378).

