## Supplementary Information for "The pyrethroid insecticide deltamethrin disrupts neuropeptide and monoamine signaling pathways in the gastrointestinal tract"

[2]Aligning Science Across Parkinson's (ASAP) Collaborative Research Network; Chevy Chase MD 20815

[3]Gangarosa Dept of Environmental Health, Rollins School of Public Health; Emory University; Atlanta GA 30329

#### Supplementary Information

Supplementary Figure S1
Supplementary Figure S2
Supplementary Table S1

### Supplementary Figure S1

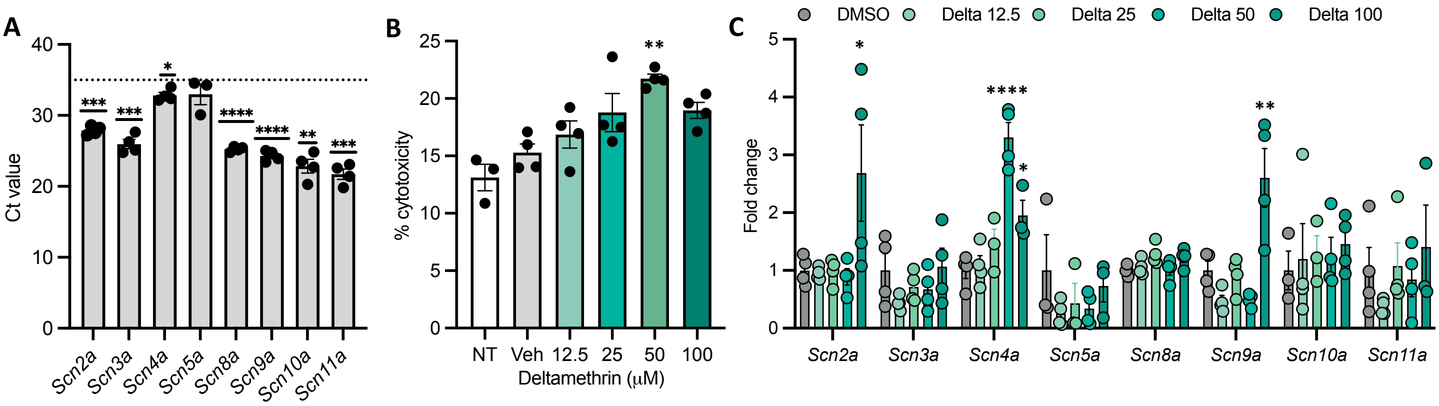


**Supplementary Fig. S1. STC-1 cells express deltamethrin-sensitive voltage-gated sodium channels. A** Ct values of each voltage-gated sodium channel (VGSC) subtype expressed in STC-1 cells. **B** Quantification of % cytotoxicity by lactate dehydrogenase (LDH) detection assay from STC-1 cells treated with 0, 12.5, 25, 50, or 100 $\mu$M deltamethrin for 24h. **C** Fold change of different VGSC subtypes from STC-1 cells treated with 0, 12.5, 25, 50, or 100 $\mu$M deltamethrin for 24h, as determined by qPCR. **A-C** All data points represent averages of technical duplicates from an individual well (n = 3-4). Data are depicted as mean $\pm$ SEM and compared by ordinary one-way ANOVA with Dunnett's multiple comparisons tests (**B, C**), one-sample t-tests against a Ct cutoff value of 35 (**A**), represented by a dashed line. *p<0.05, **p< 0.005, ****p<0.0001.

### Supplementary Figure S2

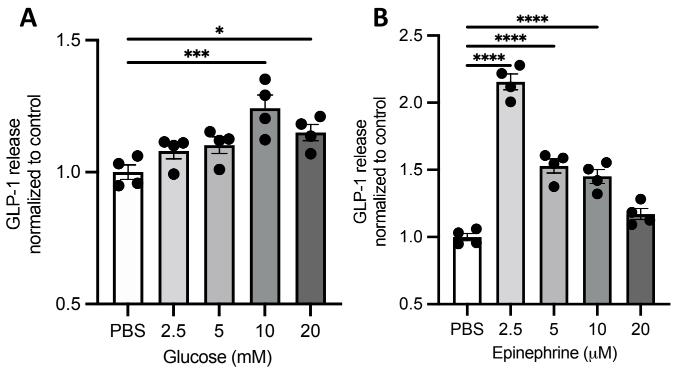


**Supplementary Fig. S2. Glucose and epinephrine evoke GLP-1 release in STC-2 cells. A** Quantification of GLP-1 release by ELISA from STC-1 cells treated with 0, 2.5, 5, 10, or 20 mM glucose for 1h. **B** Quantification of GLP-1 release by ELISA from STC-1 cells after treatment with 0, 2.5, 5, 10, or 20 $\mu$M epinephrine for 1h. **A-B** All data points represent averages of technical duplicates from individual samples where n = 3-4 per group. Data are depicted as mean $\pm$ SEM and compared by ordinary one-way ANOVA with Dunnett's multiple comparisons test comparing each group to the control (**A-B**). *p<0.05, ***p<0.001, ****p<0.0001.

#### Supplementary Table 1

| RESOURCE TYPE | RESOURCE NAME | SOURCE | IDENTIFIER | NEW/REUSE |
|---|---|---|---|---|
| Protocol | STC-1 cell culture | protocols.io | dx.doi.org/10.17504/protocols.io.j8nlk98kwv5r/v1 | reuse |
| Protocol | Dye-based intestinal motility | protocols.io | dx.doi.org/10.17504/protocols.io.eq2ly6pjwgx9/v1 | new |
| Protocol | Total GI transit assay | protocols.io | dx.doi.org/10.17504/protocols.io.14egn9676l5d/v1 | reuse |
| Protocol | Fecal Output | protocols.io | dx.doi.org/10.17504/protocols.io.rm7vzj3j5lx1/v1 | reuse |
| Protocol | LDH cytotoxicity assay | protocols.io | dx.doi.org/10.17504/protocols.io.261ger51yl47/v1 | reuse |
| Protocol | GLP-1 ELISA assay | protocols.io | dx.doi.org/10.17504/protocols.io.j8nlkokm6v5r/v1 | reuse |
| Protocol | Western blot | protocols.io | dx.doi.org/10.17504/protocols.io.kxygxwyxkv8j/v1 | reuse |
| Protocol | RNA extraction | protocols.io | dx.doi.org/10.17504/protocols.io.e6nvwdyn7lmk/v1 | reuse |
| Protocol | cDNA synthesis | protocols.io | dx.doi.org/10.17504/protocols.io.14egn9676l5d/v1 | reuse |
| Protocol | RT qPCR | protocols.io | dx.doi.org/10.17504/protocols.io.36wgqdn1ovk5/v1 | new |
| Experimental model: Cell line | STC-1 cell line | ATCC | Cat#: CRL-3254 | reuse |
| Experimental model: Organism/strain | C57BL/6J mice | Jax | RRID: IMSR_JAX:000664 | reuse |
| Chemical, peptide, or recombinant protein | Deltamethrin | Chem Service | Cat#: N-11579-250MG | reuse |
| Chemical, peptide, or recombinant protein | Glucose | Sigma | Cat#: G8270-1KG | reuse |
| Chemical, peptide, or recombinant protein | Epinephrine | Sigma | Cat#: E4250-1G | reuse |

| | | | | |
|---|---|---|---|---|
| Chemical, peptide, or recombinant protein | Carmine red dye | Sigma | Cat#: C1022 | reuse |
| Chemical, peptide, or recombinant protein | Ensure Original mixed-meal nutrient drink | Abbott Pharmaceuticals | SKU#: 57243 | reuse |
| Critical commercial assay | GLP-1 ELISA kit | Millipore | Cat#: EGLP-35K | reuse |
| Critical commercial assay | LDH cytotoxicity kit | ProMega | Cat#: G1780 | reuse |
| Critical commercial assay | U-PLEX Metabolic Hormones Combo 1 for mouse | MSD | Cat#: K15306K-2 | reuse |
| Oligonucleotide | *Tph1* qPCR primer forward | IDT | CCATCTTCCGAGAGCTAAACAAA | new |
| Oligonucleotide | *Tph1* qPCR primer reverse | IDT | TCTTCCCGATAGCCACAGTATT | new |
| Oligonucleotide | *Tph2* qPCR primer forward | IDT | TCGAAATCTTCGTGGACTGCG | new |
| Oligonucleotide | *Tph2* qPCR primer reverse | IDT | CGGATTCAGGGTCACAATGGT | new |
| Oligonucleotide | *Vmat1* qPCR primer forward | IDT | GTCCCGGAAGCTGGTGTTG | new |
| Oligonucleotide | *Vmat1* qPCR primer reverse | IDT | ACAGTGAGCAGCATATTGTCC | new |
| Oligonucleotide | *Vmat2* qPCR primer forward | IDT | CGCAAGCTGATCCTGTTCATC | new |
| Oligonucleotide | *Vmat2* qPCR primer reverse | IDT | ACGACGGTGAGCAGCATGT | new |
| Oligonucleotide | *Slc6a4* qPCR primer forward | IDT | TATCCAATGGGTACTCCGCAG | new |
| Oligonucleotide | *Slc6a4* qPCR primer reverse | IDT | CCGTTCCCCTTGGTGAATCT | new |
| Oligonucleotide | *Ddc* qPCR primer forward | IDT | TAGCTGACTATCTGGATGGCAT | new |
| Oligonucleotide | *Ddc* qPCR primer reverse | IDT | GTCCTCGTATGTTTCTGGCTC | new |
| Oligonucleotide | *Maoa* qPCR primer forward | IDT | GTGAATGTCAATGAGCGTCTAGT | new |
| Oligonucleotide | *Maoa* qPCR primer reverse | IDT | TCAACAGGGATCTCTTTTCCCA | new |
| Oligonucleotide | *Comt* qPCR primer forward | IDT | CTGGGGGTTGGTGGCTATTG | new |
| Oligonucleotide | *Comt* qPCR primer reverse | IDT | CCCACTCCTTCTCTGAGCAG | new |

| | | | | |
|---|---|---|---|---|
| Oligonucleotide | *Glp1r* qPCR primer forward | IDT | TCAGAGACGG TGCAGAAATGG | new |
| Oligonucleotide | *Glp1r* qPCR primer reverse | IDT | ATCAAAGGTCC GGTTGCAGAA | new |
| Oligonucleotide | *Gapdh* qPCR primer forward | IDT | TGGCCTTCCGT GTTCCTA | new |
| Oligonucleotide | *Gapdh* qPCR primer reverse | IDT | GAGTTGCTGTT GAAGTCGCA | new |
